# Aggressive Neuroblastomas Start Growing after Infancy

**DOI:** 10.64898/2025.11.28.691194

**Authors:** Daniel L. Monyak, Shannon T. Holloway, Graham J. Gumbert, Kellie Kim, Allison Fong, Lars J. Grimm, Jeffrey R. Marks, Darryl Shibata, Marc D. Ryser

**Affiliations:** Trinity College of Arts and Sciences, Duke University; Durham, NC, USA; Department of Population Health Sciences, Duke University School of Medicine; Durham, NC, USA; Department of Pathology, University of Southern California Keck School of Medicine; Los Angeles, CA, USA; Department of Radiology, Duke University School of Medicine; Durham, NC, USA; Department of Surgery, Duke University School of Medicine; Durham, NC, USA; Department of Mathematics, Duke University; Durham, NC, USA; Faculty of Medicine, University of Geneva; Geneva, Switzerland

## Abstract

In neuroblastoma, population screening during infancy failed to lower mortality because it primarily detected biologically indolent tumors rather than aggressive, life-threatening disease. The failure may reflect limited screening sensitivity, or disease onset after infancy among aggressive tumors. Here, we used an epigenetic mitotic clock based on fluctuating CpG DNA methylation to estimate patient-specific tumor mitotic ages and calendar ages in a cohort of unscreened children diagnosed with neuroblastoma. Aggressive cancers (stage 4) primarily started growing after the first year of life, making them undetectable by screening during infancy. In contrast, biologically more indolent tumors (stages 1, 2, 3 and 4S) often started growing in utero or during the first year of life, and affected children had better survival outcomes. Due to a short preclinical detectable phase of aggressive neuroblastomas, reducing mortality through screening is impractical as it would require frequent screening among older children. Patient-specific tumor-age estimation may help refine screening windows and improve early-detection strategies in other cancers where screening has so far failed to yield substantial mortality reductions.

## INTRODUCTION

Early detection is a cornerstone of cancer control, aiming to identify tumors before clinical symptoms emerge— when they are less extensive, and more likely to be curable. The effectiveness of this approach critically depends on the sojourn time—the interval during which a malignant lesion is screen-detectable but asymptomatic. Screening tests preferentially uncover slow-growing tumors with long sojourn times; rapidly growing tumors are more difficult to intercept because their sojourn times are often short compared to the screening interval. Due to this length-time bias,^1^ screening can only incur a mortality reduction if the sojourn times of aggressive cancers are sufficiently long for effective interception. If this is not the case, screening instead leads to overdiagnosis and overtreatment in the absence of a mortality benefit.^2^

In the case of neuroblastoma, a common pediatric cancer, substantial efforts were made in the 1990s to develop a urine-based early detection screening test for cancer catecholamine metabolites.^3^ Screening in the first year of life increased the incidence of biologically indolent neuroblastoma, without reducing advanced-stage disease.^4,5^ Mortality was unaffected by screening during infancy, and population-based neuroblastoma screening was eventually abandoned altogether.^3,6-8^

Neuroblastomas can begin to grow before birth,^9,10^ and mutations found in tumors at the time of removal have been inferred to occur as early as the first trimester.^11,12^ The failure of screening suggests two possibilities: either the test did not manage to detect asymptomatic aggressive tumors during the first year of life, or these aggressive tumors simply had not yet begun to grow during infancy **(Figure 1A**). To identify the more likely reason of failure, we developed an epigenetic clock to determine the age of neuroblastoma tumors based on fluctuating CpG (fCpG) DNA methylation (**Figure 1B**). Disease-site tailored versions of this clock have previously been used to characterize the temporal landscape of breast cancers,^13^ track the evolution of hematologic malignancies,^14^ and quantify the dynamics of intestinal crypts.^15^

**Figure 1.**
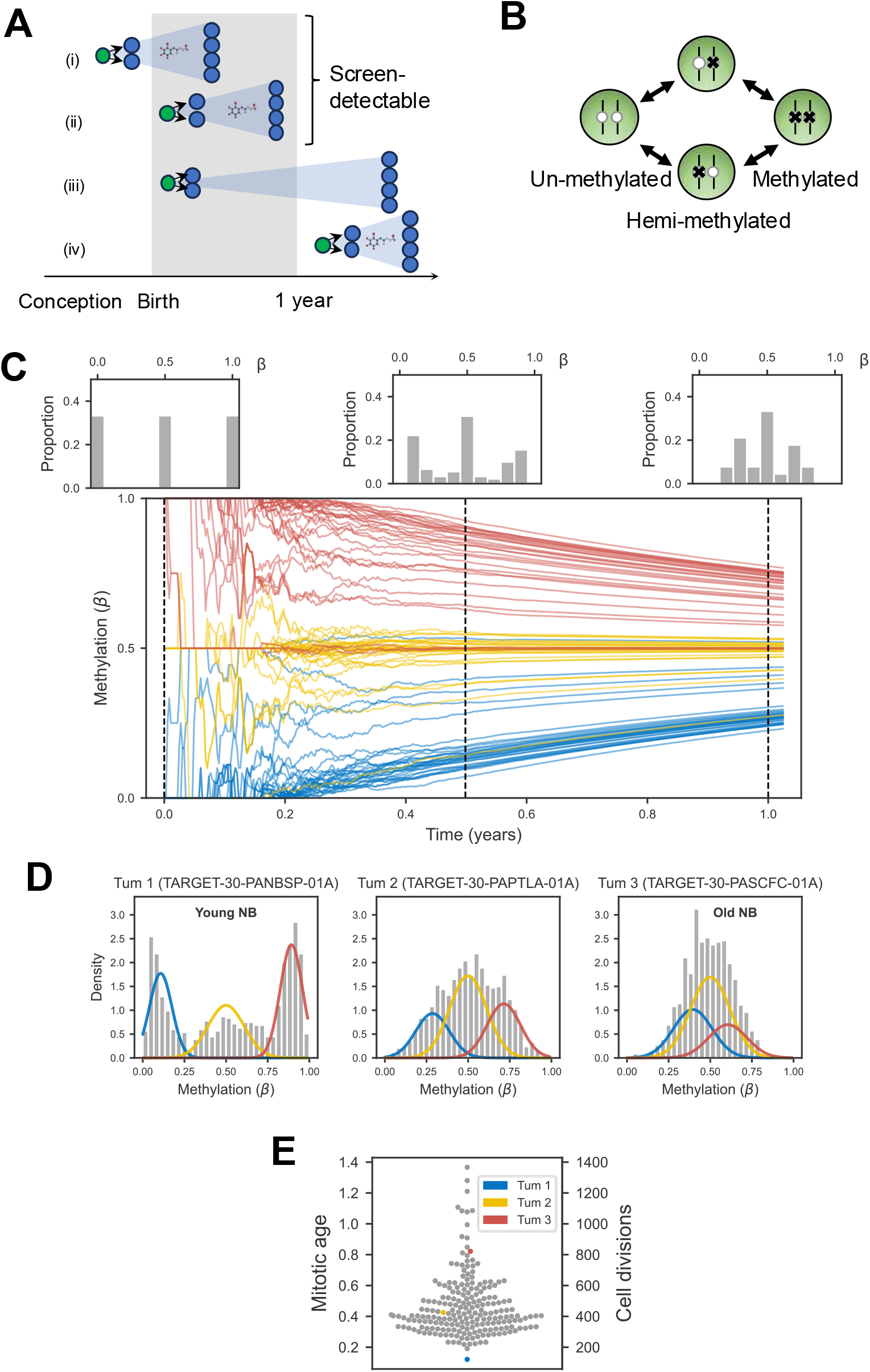
Inference of tumor mitotic age from fCpG methylation dynamics. **(A)** Neuroblastoma screening (by urinary catecholamines test) during infancy can detect tumors that started growing (green circle) before (i) or shortly after (ii) birth. Screening fails if the tumor does not produce measurable concentrations of urinary catecholamines (iii) or if the tumor starts growing after the screening period (iv). **(B)** Fluctuating CpG (fCpG) sites stochastically oscillate between the unmethylated, hemi-methylated, and methylated states. **(C)** In the most recent common ancestor cell of a growing neuroblastoma tumor (time=0), individual fCpGs are unmethylated (blue), hemi-methylated (yellow), or methylated (red). Over time, the average methylation value of either initial state converges to 50% methylation. Measuring the methylation distribution across an ensemble of fCpG sites (see histograms at 6 and 12 months) provides a measure of tumor mitotic age: younger tumors have a more dispersed methylation distribution whereas older tumors have a more concentrated methylation distribution. **(D)** Examples of fCpG methylation distributions for younger (Tum 1), middle aged (Tum 2), and older (Tum 3) neuroblastomas in the TARGET cohort. The solid curves represent the inferred distributions for fCpG sites that started at 0 (blue), 50% (yellow) or 100% (red) methylation in the progenitor cell. **(E)** Estimated (normalized) tumor mitotic ages (left axis) of all neuroblastomas in the TARGET cohort (N = 213); conversion to number of cell divisions (right axis) is based on an assumed (de-)methylation rate of 10^−3^.

Using this clock, we estimated the mitotic age of individual tumors—that is the number of cell doublings between the most common recent ancestor cell and the resected specimen—in an unscreened cohort of NB patients. Aggressive neuroblastomas are mitotically younger than biologically indolent tumors and predominantly initiated growth after the first year of life. Short sojourn times rather than low test sensitivity is thus most likely what undermined the effectiveness of screening during infancy to reduce neuroblastoma mortality.

## RESULTS

### Estimating Tumor Mitotic Age in Neuroblastoma Patients

We developed a neuroblastoma-specific epigenetic clock following our previously described approach.^13^ In brief, we analyzed DNA methylation (850K EPIC‘ array) from 213 tumors from the neuroblastoma TARGET cohort^16^ to identify a set of 1,000 fluctuating CpG sites (**Supplementary Data S1**) whose average methylation values, or *β*-values (between 0 and 1), were balanced (**Supplementary Figure 1A**), did not correlate with patient age (**Supplementary Figure 1B**) and had large inter-tumor variability (**Supplementary Figure 1C**).

The fCpG clock is well suited to infer tumor mitotic ages because it resets at the start of growth. Indeed, fCpG methylation in the most recent common ancestor cell is either *β* = 0 (unmethylated site), *β* = 0.5 (hemi-methylated site), or *β* = 1 (methylated site). As the tumor grows through cell proliferation, the average methylation at fCpG sites that were originally unmethylated (*β* = 0) or fully methylated (*β* = 1) will slowly drift to *β* = 0.5 (**Figure 1C)**. Therefore, by measuring the methylation distribution in an ensemble of fCpGs, or clock set, we can distinguish younger tumors with discrete peaks in the methylation distribution (low epigenetic entropy) from older tumors with unimodal distributions centered around *β* = 0.5 (high epigenetic entropy; **Figure 1D**).

To infer tumor mitotic age from a clock set of fCpG sites, we estimated the location of the left and right peaks in the *β*-value histograms (**Figure 1D**, blue and red curves) and converted peak location to tumor mitotic age based on a stochastic model describing the methylation dynamics of fCpG sites; see **Methods** for details. In the TARGET cohort, the median tumor mitotic age was 400 generations, ranging from 121 to 1,367 generations (**Figure 1E**).

### Aggressive Neuroblastomas Start Growing after Infancy

The conversion from tumor mitotic age to tumor calendar age requires calibration of the molecular clock. In adult malignancies, the unobservable start of expansion from a single progenitor cell can predate the date of diagnosis by decades, rendering direct calibration challenging.^13^ In pediatric cancers, however, tumor calendar age is more constrained because the start of growth cannot predate conception. Noting that neuroblastomas can be physically detected as early as the first trimester,^9,10^ we used patients who were diagnosed shortly after birth to convert tumor mitotic age (cell doublings) into tumor calendar ages (years); see **Methods** for details.

As visualized in **Figure 2**, most neuroblastomas diagnosed during infancy (first year of life) started growing in utero or shortly after birth. In 166/167 (99.4%) of cancers diagnosed after 1.5 years of patient age, tumor growth started after the first year of patient age, that is after the screening period used in earlier trial. Overall, 163 of 167 stage 4 tumors (97.6%) started to grow after one year of age, whereas 42 of 46 (91.3%) of biologically more indolent tumors (stages 1,2,3,4S) started to grow before one year of age.

**Figure 2.**
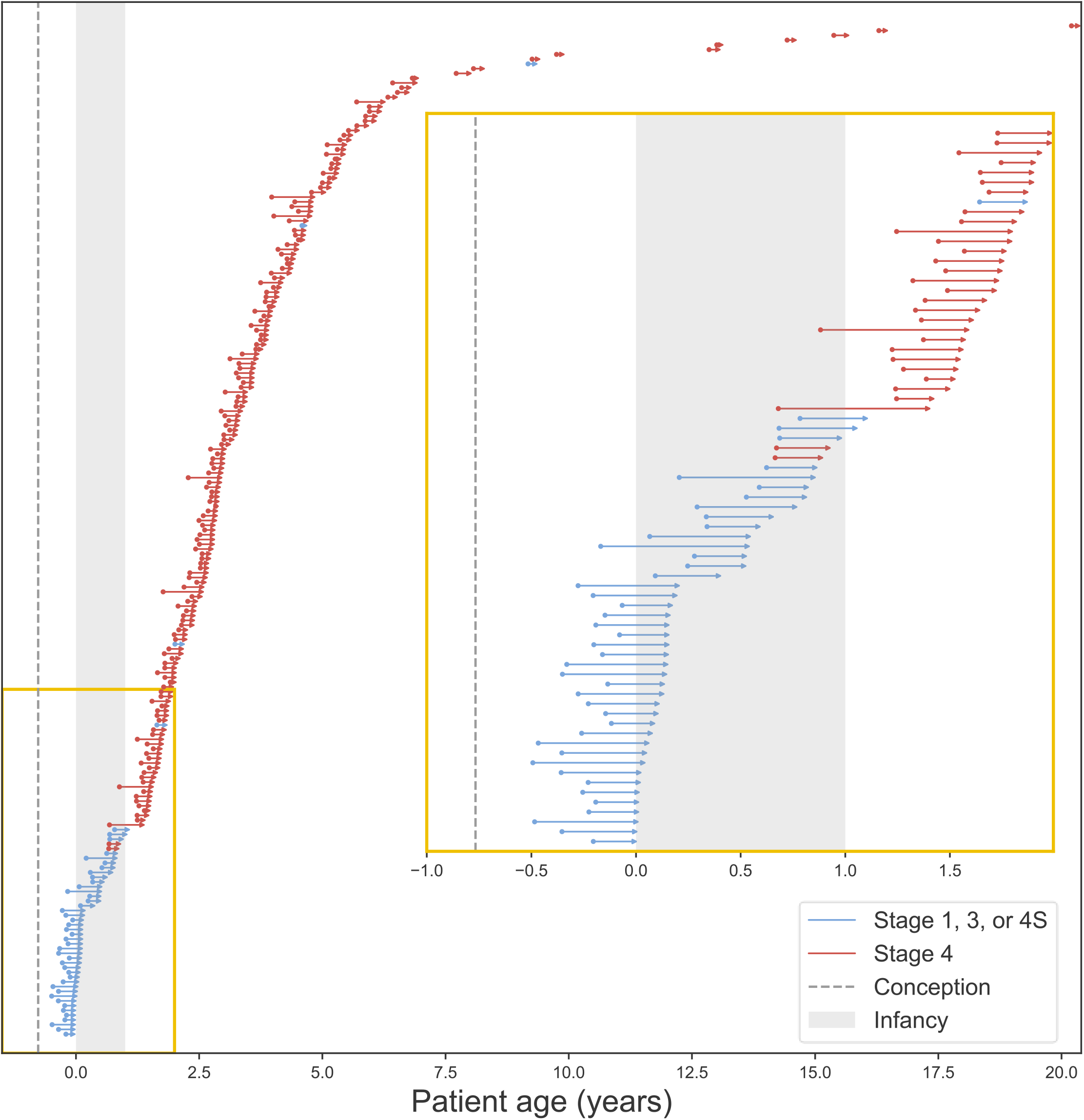
Tumor calendar ages. For each neuroblastoma tumor in the TARGET cohort, the estimated tumor calendar age is represented by a horizontal line that connects the start of its expansion (left dot) and its detection (right arrow). The dotted vertical line represents the time of conception, and the shaded gray interval spans first year of patient life. Insert: magnified view of tumors discovered in patients diagnosed before 2.5 years of age.

The epigenetic clock-based tumor age estimate includes the initial expansion phase when the growing mass is too small to be screen-detectable. Nevertheless, tumor calendar age provides an upper bound to the sojourn time—the period when the tumor is asymptomatic but screen-detectable. Based on our analysis, the upper bound to the mean sojourn time among aggressive stage 4 tumors is 109 days (range, 29-329 days), compared to 126 days (range, 46-262 days) among indolent (stages 1,2,3,4s) tumors (P = 0.03; two-sided t-test).

In summary, in this cohort, even the most sensitive screening test would have failed to detect a clinically significant proportion of lethal neuroblastomas during infancy. These findings suggest that the historical failure of neuroblastoma screening to reduce mortality^3,8^ may reflect the fact that aggressive tumors typically started growing after the screening test was administered.

### Tumor Mitotic Age Correlates with Prognostic Factors

Among unscreened patients, neuroblastoma usually presents as a palpable mass. By the time they reach a macroscopic size, slow-growing tumors are expected to have accumulated more cell divisions and appear mitotically older compared to fast-growing and aggressive tumors. We tested this hypothesis by investigating the relationship between tumor mitotic age and prognostic factors in an unscreened cohort of children diagnosed with neuroblastoma.

As predicted, stage 1 and 4S tumors were mitotically older compared to metastatic stage 4 cancers (**Figure 3A**). Similarly, tumors classified as high-risk were mitotically younger tumors compared to low-risk tumors (**Figure 3B**). Tumors with *MYCN* amplification (a marker of aggressiveness) were mitotically younger than tumors without *MYCN* amplification (**Figure 3C**), and patients diagnosed before the age of 1.5 years had mitotically older tumors (**Figure 3D**). We validated these finding in an independent cohort of 105 clinically diagnosed neuroblastoma tumors (**Supplementary Figure S2**).^17^

**Figure 3.**
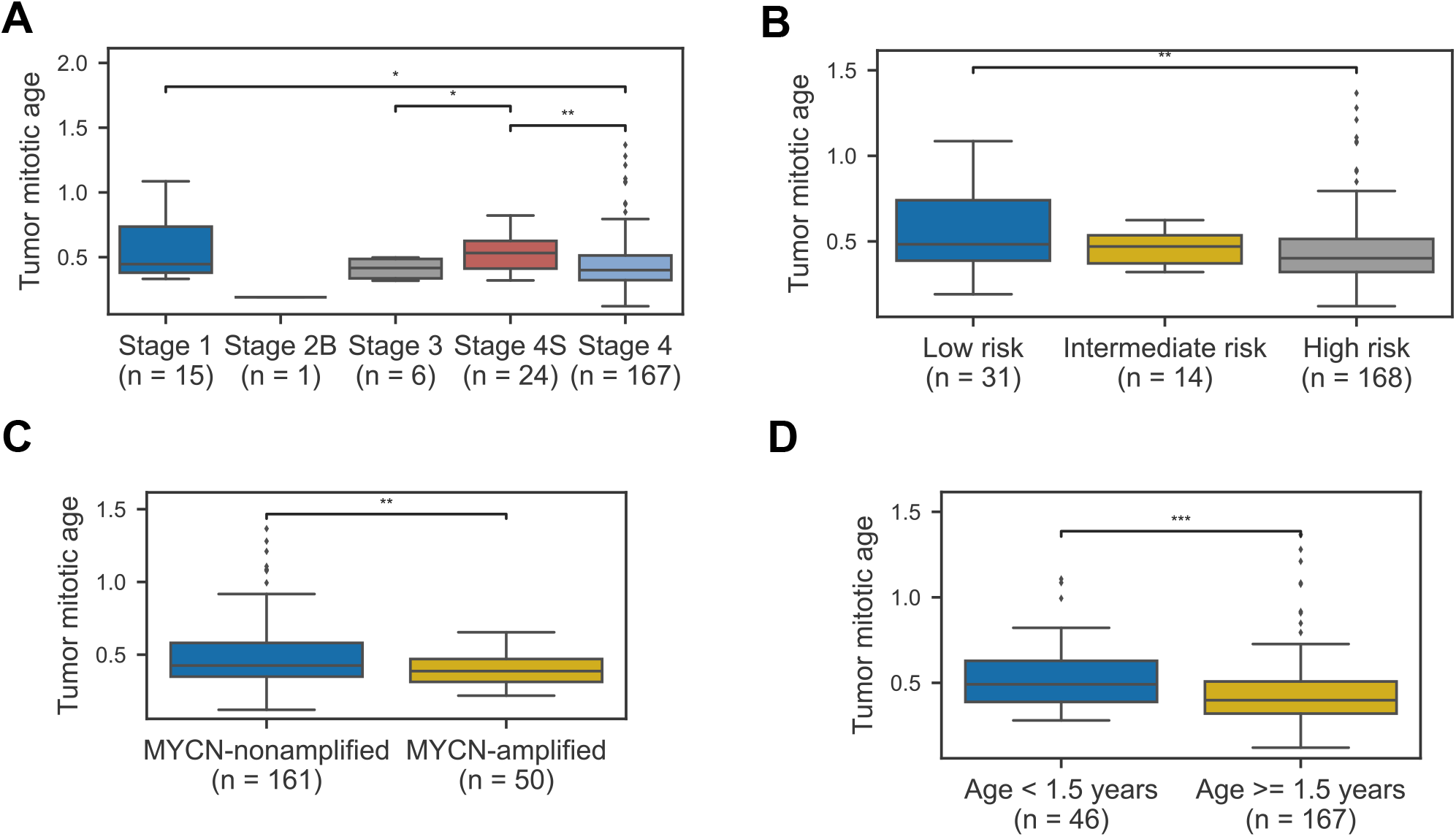
Tumor mitotic age vs. clinicopathological variables. The distribution of tumor mitotic age in the TARGET cohort, by tumor stage **(A)**; risk category **(B)**; *MYCN* amplification status **(C);** and patient age, stratified at 1.5 years **(D)**. Pairwise comparisons of medians were performed using a two-sided Wilcoxon rank-sum test (*P< 0.05, **P< 0.01, ***P< 0.001). Similar findings were observed in an independent cohort (see Supplemental Fig. S3).

Finally, to determine molecular correlates, we combined the mitotic age estimates with gene expression data (RNA-seq) to perform a gene-set enrichment analysis (GSEA). Mitotically younger neuroblastomas were enriched for proliferation pathways whereas no pathway enrichment was found in older tumors (**Supplementary Figure S3**). Thus, consistent with their more aggressive biology, the fCpG clock infers that more deadly neuroblastomas grow faster.

### Tumor Age Predicts Survival

We next examined the relationship between tumor mitotic age and survival. In a univariate Cox model, younger tumor mitotic age was associated with lower overall survival (HR 0.16; 95% CI, 0.046-0.55; P = 0.004; **Figure 4A**). When considering patients diagnosed with stage 4 cancers only, younger mitotic age remained a predictor of poor survival (HR 0.26; 95% CI, 0.079-0.84; P = 0.025; **Figure 4B**). We note that in pediatric cancers, overall survival over a medium-term follow-up is a good proxy for disease-specific survival.

**Figure 4.**
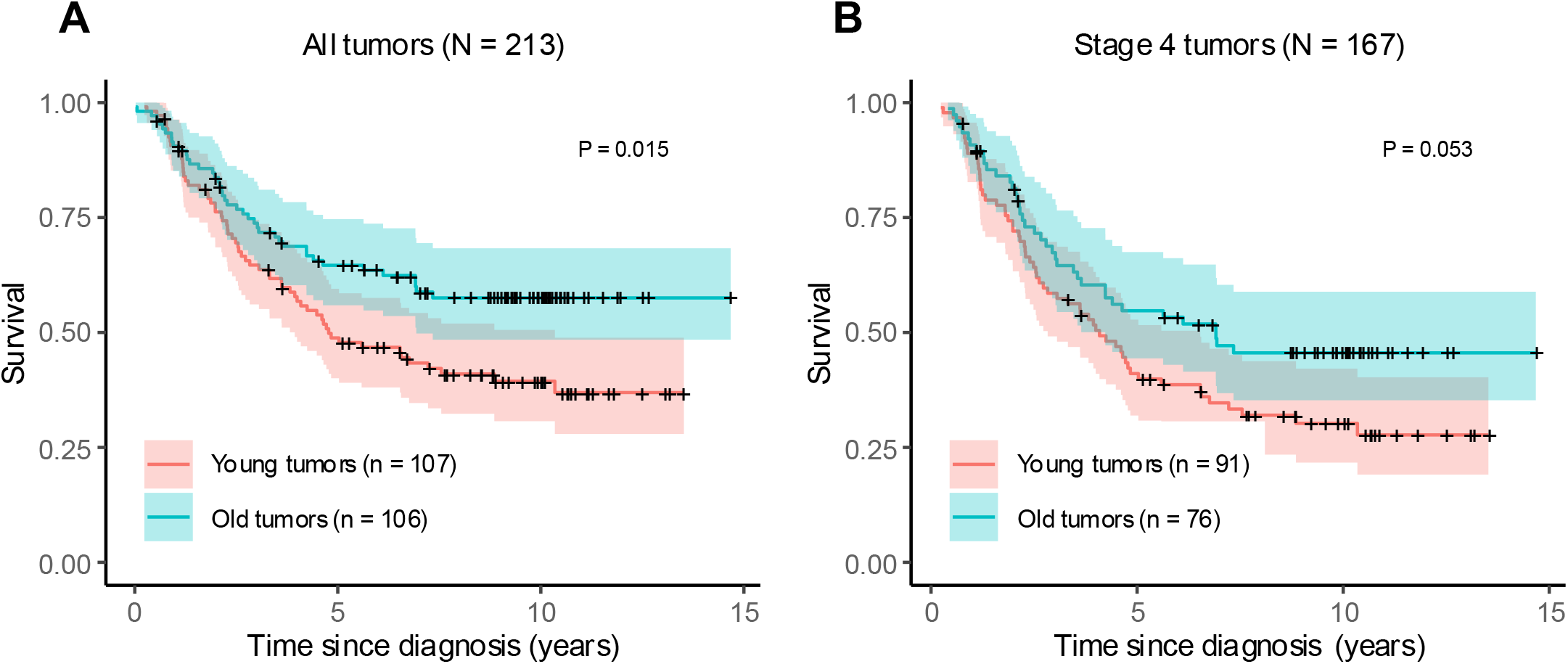
Tumor mitotic age vs. survival. Kaplan-Meier curves for overall survival in the TARGET cohort, stratified by tumor mitotic age (young: below median; old: above median). Results are shown for all tumors **(A)** and stage 4 tumors only **(B)**. Survival was compared between old and young tumors using the log-rank test. See main text for results from a Cox proportional hazards model using continuous mitotic tumor age as predictor.

## DISCUSSION

Neuroblastoma screening increased the detection and treatment biologically favorable tumors without reducing the incidence of lethal cancers and mortality.^3,8^ What remains unclear, however, is why lethal cancers were missed in the first place: was the test’s sensitivity to detect aggressive phenotypes too low, or did these tumors only start growing after the screening period? Applying an epigenetic molecular clock to a cohort of unscreened neuroblastoma patients, we show that most aggressive stage 4 tumors started growing after the first year of life. As such their tumors were not present—and thus not detectable—during infancy. In contrast, most tumors with a more favorable biology started growing before or shortly after birth, rendering them a priori screen-detectable during infancy. Our findings suggest that the failure of neuroblastoma screening was due to a mismatch between screening window and onset of lethal disease, rather than insufficient sensitivity of the test.

These findings raise the question whether extending screening beyond the first year of life could provide a way to improve outcomes in patients affected by neuroblastoma. In our analysis, we estimated the sojourn times of lethal stage 4 cancers to be very short—of the order of at most a few months—suggesting a very limited opportunity for early detection, even under very frequent screening after infancy.

An increase in the incidence of favorable tumors is a common occurrence in cancer screening and has also been documented in cancers of the breast,^18^ prostate^19^ and lung.^20^ However, for these cancer types, the overdiagnosis of good-outcome cancers is accompanied by the detection of a proportion of aggressive cancers which in turn produces a mortality benefit for the screening test. The situation in neuroblastoma screening is different in that the incidence of lethal cancers remained unchanged,^7,21^ resulting in very high rates of overdiagnosis in the absence of any screening benefit.

This work highlights how tumor-level age estimates from fCpG molecular clocks can inform and guide screening practices. Methylation-based molecular clocks were originally introduced to measure biological tissue aging,^22-25^ and have been successfully applied to quantifying cancer risk.^26,27^ A common feature of many of these clocks is that they leverage a unidirectional drift in methylation of tissue over time.^27^ Our clock differs in that it starts with— or is reset to—the fCpG methylation pattern in the most recent common ancestor cell. This reset, along with rapid, balanced methylation and demethylation at fCpG sites, allows us to rank cancers by their mitotic age, which may vary by only a few cell divisions.

Calibrating a cancer clock—that is converting mitotic to calendar age—is challenging because the calendar ages of tumors are invariably unknown. In the case of pediatric cancers like neuroblastoma, however, the natural age constraint by the date of conception provides an opportunity for calibration. The type of molecular clock we adapted here to the neuroblastoma setting had previously been used to study stem cell turnover in normal human tissues,^28^ to show that acute leukemias exhibit synchronized “W-shaped” fCpG distributions, while chronic leukemias display desynchronized, unimodal distributions,^28^ and to characterize the temporal landscape of human breast cancer.^13^ Like for neuroblastomas, breast cancer mitotic ages vary, with more aggressive tumors tending to be mitotically younger.

Our study has several limitations. First, our analyses are inferential in nature, and the absence of a direct empirical measure of tumor age precludes a gold-standard validation of our findings. However, our mitotic age estimates align well with established neuroblastoma biology, prognostic factors and survival outcomes. Furthermore, the approach has been thoroughly tested in the setting of breast cancer^13^ and hematologic malignancies.^14^ Second, while conception provides an upper bound for tumor calendar age, our calibration based on the youngest patients may not be applicable to older children if methylation fluctuation rates change as a function of the child’s age. Third, the epigenetic clock measures only the age of the final clonal expansion. Earlier precursor cells that predate this expansion, and whose existence can be inferred from copy-number alterations and somatic mutations,^11,12^ are not captured by this approach. However, we note that small, isolated patches of precursor cells are not detectable by screening either.

Cancer early detection efforts crucially depend on organ-specific natural history of disease. To date, these dynamics have primarily been characterized by summary statistics such as the mean sojourn time, as obtained by fitting mathematical models to screened cohorts. The approach used in this work provides an opportunity to move from statistical averages to tumor-specific mitotic ages and sojourn times. Drawing from readily available methylation array data, the molecular clock we used in this work can be further generalized to estimate tumor-specific mitotic ages and sojourn times across various cancer types.

## ACKNOWLEDGMENTS

This work was partially supported by grant R01-CA271237 from the National Institutes of Health.

## DATA AVAILABILITY

All data used in this work are publicly available under https://portal.gdc.cancer.gov (TARGET data) and under the Gene Expression Omnibus accession number GSE73515 (German Neuroblastoma Trial). A list of the 500 fCpG sites used to estimate tumor mitotic age is provided in Supplementary Data 1.

## CODE AVAILABILITY

All Python (version 3.9.19) and R (version 4.3) code used to produce the results in this paper are found on GitHub at https://github.com/danmonyak/EpiClockNBL (MIT License).

## AUTHOR CONTRIBUTIONS

DLM, DS and MDR conceived the study. MDR and LJG secured funding. MDR, DLM, KK, AF, and STH developed and optimized the analytical methods. DLM, GG, and STH curated the datasets, maintained the code base, and performed the computational analyses. DLM prepared the figures and illustrations. All authors contributed to the interpretation of the results. MDR, DS, and JRM supervised the project. DLM, DS, and MDR drafted the manuscript, and all authors revised it critically and approved the final version.

## METHODS

### Samples

Of the 842 neuroblastomas from patients in the Therapeutically Applicable Research to Generate Effective Treatments (TARGET) program,^16^ 213 had available methylation array data (Infinium HumanMethylation450 BeadChip, Illumina, San Diego, CA, USA), which were used to select an ensemble of 1,000 fCpG sites. For a single tumor in this set, two methylation measurements were initially present but one was removed. The following variables were retrieved: patient age at diagnosis, International Neuroblastoma Staging System (INSS) stage, Children’s Oncology Group (COG) risk category, and *MYCN* amplification status. Among the 213 tumors, gene expression quantification (RNA-seq) data was available for 132. All clinical and sequencing data were retrieved from the Genomic Data Commons (GDC; https://gdc.cancer.gov) using the R package *TCGAbiolinks* (version 2.25.3). We further retrieved publicly available methylation array data (Infinium HumanMethylation450 BeadChip) from 105 neuroblastomas^17^ from patients enrolled in the German Neuroblastoma Trial (GSE73515). The following variables were retrieved: patient age group at diagnosis; INSS stage; COG risk category, and *MYCN* amplification status.

### Selection of fluctuating CpG (fCpG) sites

As previously described,^13^ we first identified CpG sites with balanced methylation and de-methylation rates, defined as having an average methylation content (*β*-value) between 0.4 and 0.6 in the TARGET cohort (***Supplementary Figure 1A***); CpG sites who were missing in ≥20 patients were excluded. In a second step, we eliminated age-confounded sites by calculating the Spearman correlation (*ρ*) between patient age at diagnosis and CpG *β*-value and eliminating sites with |*ρ*| > 0.2 (***Supplementary Figure 1B***). In a third step, the remaining CpG sites were ranked by their *β*-value variance across all tumors in the cohort and the 25% least variable sites were eliminated (***Supplementary Figure 1C***). In a final step, we selected non-clustering CpG sites. In brief, we removed sites that were correlated with each other, possibly due to the existence of latent tumor subgroups (e.g., molecular subtypes) that induce correlations between CpG sites. To achieve this, we performed K-Means clustering on the remaining CpG sites and identified the final clock set of 1,000 fCpG sites that exhibited minimal *β*-value variation between clusters; see **Supplementary Methods** for details.

### Estimation of tumor mitotic age

To estimate tumor mitotic age from the empirical *β*-value distribution, we proceeded in two steps. In the first step, we decomposed each tumor’s *β*-value distribution into three peaks of fCpG sites (see, e.g., ***Figure 1D***): the originally unmethylated fCpG sites (left peak), the originally hemi-methylated fCpG sites (middle peak), and the originally methylated fCpG sites (right peak). This was achieved by fitting a Gaussian mixture model to the empirical data using the Bayesian sampling algorithm *Stan* as (see *Gaussian mixture fitting* below) and resulted in the location of the left (*X*) and right (1 − *X*) methylation peaks. In the second step, as previously described,^13^ we converted the peak location *X* to a (normalized) tumor mitotic age *ϕ* based on a stochastic model of the fCpG methylation dynamics, *ϕ* = −log (1 − 2*X*)/2. To convert mitotic age *ϕ* to the number of cell doublings, the former is rescaled as *ϕ*/*μ*, where *μ* is the (de-)methylation probability per allele per cell division; for illustrative purposes we assumed *μ* = 10^−3^.^13^

### Gaussian mixture fitting

Each tumor’s *β*-value distribution was modeled as a mixture of three normal distributions: a left component centered at a free parameter *X*, a middle component centered at 0.5, and a right component centered at 1 − *X* (by symmetry). The proportions of the three components were denoted *ψ*_1_ (left component), *ψ*_2_ (middle component), and *ψ*_3_ = 1 − *ψ*_2_ − *ψ*_3_ (right component). The complete mixture model and corresponding data likelihood are detailed in ***Supplementary Methods***. We then combined the likelihood with the Bayesian sampler *Stan* (R package *rstan*, version 2.36.0.9000) to perform parameter inference. A Dirichlet prior with concentration (5, 10, 5) was used on the component proportions (*ψ*_1_, *ψ*_2_, *ψ*_3_); a *Beta*(1, 4) prior on *X* was chosen to reflect the fact that the left peak moves from *β* = 0 to *β* = 0.5, and a non-informative *Gamma*(2, 0.1) prior on *n* to allow flexibility in the effective allele number. Sampling from the posterior distribution of the model parameters (*X, n, ψ*_1_, *ψ*_2_, *ψ*_3_) were performed independently for each tumor, with four chains of 4,000 iterations each (2,000 warmup). Satisfactory convergence was achieved as assessed by visual inspection of trace plots, 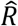 statistics (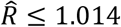 across all parameters and tumors), and effective sample sizes (ESS≥ 363 across all parameters and tumors).

### Estimation of tumor calendar age

A tumor’s (normalized) mitotic age *ϕ* is related to its calendar age *t* by *ϕ* = *tαμ*, where *α* is the cell proliferation rate. To obtain an estimate for the product *αμ* we imposed an earliest time for the start of tumor growth at 180 before birth, as suggested by previous studies that found that neuroblastomas can be detected as early as the first trimester.^11,12^

### Pathway enrichment analyses

For the TARGET cohort, we performed a gene set enrichment analysis (GSEA) using the software package *GSEA*^*29*^ to identify *Hallmark* gene sets that are correlated with tumor mitotic age *ϕ*. The analysis was performed using the Pearson correlation to rank individual genes; phenotype-permutation-based *P*-values and false-discovery rate (FDR) *Q*-values were computed using 1,000 permutations. All other inputs were kept at their defaults.

### In silico simulation of fCpG evolution

For illustration purposes, we modeled a set of 90 fCpGs in a growing tumor. We assumed an even distribution of unmethylated, hemi-methylated, and methylated initial states (n = 30 each) in the tumor’s founding, or most common recent ancestor cell. To simulate the birth death process of tumor growth, we performed time stepping (Δ*t*= 1 day), allowing each cell to divide with probability *α* to produce two daughter cells, or to die with probability *λ*. During cell division, we modeled the error rate for fCpG site methylation by allowing each allele to methylate /demethylate with equal probability *μ*. For the simulation in ***Figure 1C***, we used *α* = 0.17 and *λ* = 0.112, resulting in a tumor size of roughly 10^9^ cells after 1 year of tumor growth, and *μ* = 0.0075.

### Statistical analyses

Correlations between two continuous variables were calculated using the Pearson correlation coefficient. The medians of continuous variables were compared using a two-sided Wilcoxon rank-sum test at significance level of 0.05. For each variable, tumors with missing values of that variable were excluded. All analyses and visualizations were performed in Python (3.9.19) and R (version 4.3).

## REFERENCES

1 Duffy, S. W. et al. Correcting for Lead Time and Length Bias in Estimating the Effect of Screen Detection on Cancer Survival. American Journal of Epidemiology 168, 98–104 (2008). 10.1093/aje/kwn120

2 Srivastava, S. et al. Cancer overdiagnosis: a biological challenge and clinical dilemma. Nature Reviews Cancer 19, 349–358 (2019). 10.1038/s41568-019-0142-8

3 Woods, W. G. Screening for Neuroblastoma Using Urinary Catecholamines: The End of the Story. JNCI Cancer Spectrum 5 (2021). 10.1093/jncics/pkab042

4 Brodeur, G. M. Spontaneous regression of neuroblastoma. Cell and Tissue Research 372, 277–286 (2018). 10.1007/s00441-017-2761-2

5 Matthay, K. K. et al. Neuroblastoma. Nature reviews Disease primers 2, 16078 (2016).

6 Woods, W. G. et al. Screening of infants and mortality due to neuroblastoma. New England Journal of Medicine 346, 1041–1046 (2002).

7 Schilling, F. H. et al. Neuroblastoma screening at one year of age. New England Journal of Medicine 346, 1047–1053 (2002).

8 Berthold, F. et al. Neuroblastoma Screening at 1 Year of Age: The Final Results of a Controlled Trial. JNCI Cancer Spectrum 5 (2021). 10.1093/jncics/pkab041

9 Acharya, S. et al. Prenatally diagnosed neuroblastoma. Cancer 80, 304–310 (1997). 10.1002/(SICI)1097-0142(19970715)80:2<304::AID-CNCR19>3.0.CO;2-Y

10 Marshall, G. M. et al. The prenatal origins of cancer. Nature Reviews Cancer 14, 277–289 (2014). 10.1038/nrc3679

11 Körber, V. et al. Neuroblastoma arises in early fetal development and its evolutionary duration predicts outcome. Nature Genetics 55, 619–630 (2023). 10.1038/s41588-023-01332-y

12 Caravagna, G. Mathematical modeling of neuroblastoma associates evolutionary patterns with outcomes. Nature Genetics 55, 530–531 (2023). 10.1038/s41588-023-01358-2

13 Monyak, D. L. et al. Mapping the temporal landscape of breast cancer using epigenetic entropy. Communications Biology 8, 1477 (2025). 10.1038/s42003-025-08867-2

14 Gabbutt, C. et al. Fluctuating DNA methylation tracks cancer evolution at clinical scale. Nature, 1–10 (2025).

15 Gabbutt, C. et al. Fluctuating methylation clocks for cell lineage tracing at high temporal resolution in human tissues. Nature biotechnology 40, 720–730 (2022).

16 Institute, N. C. TARGET: Therapeutically Applicable Research to Generate Effective Treatments. (2024). <https://www.cancer.gov/ccg/research/genome-sequencing/target>.

17 Henrich, K.-O. et al. Integrative Genome-Scale Analysis Identifies Epigenetic Mechanisms of Transcriptional Deregulation in Unfavorable Neuroblastomas. Cancer Research 76, 5523–5537 (2016). 10.1158/0008-5472.Can-15-2507

18 Ryser, M. D. & Etzioni, R. B. Estimation of Breast Cancer Overdiagnosis in a U.S. Breast Screening Cohort. Ann Intern Med 175, W116–W117 (2022). 10.7326/L22-0277

19 Etzioni, R. et al. Overdiagnosis due to prostate-specific antigen screening: lessons from US prostate cancer incidence trends. Journal of the National Cancer Institute 94, 981–990 (2002).

20 Patz, E. F. et al. Overdiagnosis in low-dose computed tomography screening for lung cancer. JAMA internal medicine 174, 269–274 (2014).

21 Woods, W. G. et al. A population-based study of the usefulness of screening for neuroblastoma. the Lancet 348, 1682–1687 (1996).

22 Hannum, G. et al. Genome-wide Methylation Profiles Reveal Quantitative Views of Human Aging Rates. Molecular Cell 49, 359–367 (2013). 10.1016/j.molcel.2012.10.016

23 Horvath, S. DNA methylation age of human tissues and cell types. Genome Biology 14, 3156 (2013). 10.1186/gb-2013-14-10-r115

24 Youn, A. & Wang, S. The MiAge Calculator: a DNA methylation-based mitotic age calculator of human tissue types. Epigenetics 13, 192–206 (2018). 10.1080/15592294.2017.1389361

25 Zhou, W. et al. DNA methylation loss in late-replicating domains is linked to mitotic cell division. Nature Genetics 50, 591–602 (2018). 10.1038/s41588-018-0073-4

26 Bell, C. G. et al. DNA methylation aging clocks: challenges and recommendations. Genome Biology 20, 249 (2019). 10.1186/s13059-019-1824-y

27 Teschendorff, A. E. & Horvath, S. Epigenetic ageing clocks: statistical methods and emerging computational challenges. Nature Reviews Genetics 26, 350–368 (2025).

28 Gabbutt, C. et al. Fluctuating methylation clocks for cell lineage tracing at high temporal resolution in human tissues. Nature Biotechnology 40, 720–730 (2022). 10.1038/s41587-021-01109-w

29 Subramanian, A. et al. Gene set enrichment analysis: A knowledge-based approach for interpreting genome-wide expression profiles. Proceedings of the National Academy of Sciences 102, 15545–15550 (2005). doi:10.1073/pnas.0506580102

