## Supplementary Methods for "Aggressive Neuroblastomas Start Growing after Infancy"

### Supplementary Methods
#### *Aggressive Neuroblastomas Start Growing after Infancy*

Monyak et al., 2026

#### 1 Selection of non-clustering fCpGs

The first three steps of the fCpG selection process are described in the main text (**Methods**). Here, we describe the fourth and final step that seeks to identify fluctuating CpG (fCpG) sites that do not exhibit clustering across tumors. Indeed, it is possible that latent tumor subgroups (e.g., molecular subtypes) induce correlations between the remaining CpG sites, although individual sites are on average balanced (i.e., $\bar{\beta} \in [0.4, 0.6]$) and dispersed (i.e., large inter-tumor variance).

To this end, we identified latent subgroups using k-means clustering and then eliminated CpG sites that showed disparate levels of methylation between subgroups. As explained in more detail below, we introduced a CpG site-specific clustering weight $w_s$ ($s \in \{s_1, ..., s_n\}$) as a normalized measure of the site's average difference in methylation level between clusters. We then selected the 1,000 CpGs that had the lowest clustering weights to identify the final clock set of fCpGs. We note that this method was also used because directly calculating the correlation between all pairs of CpG $\beta$-value vectors was far too inefficient for over 10,000 CpGs and 213 tumors.

##### Calculating the clustering weights

To calculate the CpG site-specific clustering weight $w_s$, we performed k-mean clustering on all tumors using the methylation $\beta$-values of the CpG sites $s \in \{s_1, ..., s_n\}$ as their features, with $R = 4$ clusters. We settled on $R = 4$ because $w_s$ were found to be insensitive to a further increase in the number of clusters. For cluster $i$ and site $s$, we then calculated $C_i(s)$ as the average $\beta$-value of the site in the cluster, and used the absolute difference to quantify a site-specific cluster difference $|C_i(s) - C_j(s)|$.

When comparing a CpG site between clusters, we gave less importance to clusters that contained fewer tumors by calculating a weight, $k_{ij} = n_i n_j$, for each pair of clusters $(i,\ j)$, where $n_i$ and $n_j$ are the number of tumors in the $i^{th}$ and $j^{th}$ clusters, respectively. Finally, we divided these weights by a normalization factor:

$$K = \sum_{i=1}^{R-1} \sum_{j=i+1}^{R} k_{ij}$$

The full formula for clustering weight $w_s$ is as follows:

$$w_s = \frac{1}{K} \sum_{i=1}^{R-1} \sum_{j=i+1}^{R} k_{ij} \big| C_i(s) - C_j(s) \big|.$$

### 2 Gaussian mixture model: likelihood derivation

To decompose the empirical methylation distribution of a tumor into the three peaks stemming from initially unmethylated, hemi-methylated, and methylated CpG sites, we use a Gaussian mixture model. The measured methylation value $Y_i \in [0, 1]$ at fCpG site $i$ is then modeled as

$$Y_i \sim \psi_1 \mathcal{N}(m_1, \sigma_1^2) + \psi_2 \mathcal{N}(m_2, \sigma_2^2) + \psi_3 \mathcal{N}(m_3, \sigma_3^2), \tag{1}$$

where $\psi_1 + \psi_2 + \psi_3 = 1$.

We start by parameterizing the left peak component with parameters $(m_1, \sigma_1^2)$, that is the ensemble of fCpG sites that originated in the fully unmethylated state. We denote by $n$ the (effective) number of alleles measured by the assay and denote by $S_{ij}$ the methylation status of allele $j$ at CpG site $i$. We make the simplifying assumption that the random variables $S_{ij} \sim Bernoulli(X)$, where $X$ is the true methylation value of the sample, are independent and identically distributed. Although the evolution of fCpG sites is correlated through the underlying birth-death process of the tumor cells, we note satisfactory data fits of the resulting model, providing a post hoc justification for the *i.i.d.* assumption. We note that $Y_i = \frac{1}{n} \sum_{j=1}^{n} S_{ij}$ such that, by invoking the Central Limit Theorem for $n \to \infty$

$$Y_i | \; left \sim \mathcal{N}\left(\mathbb{E}(S_{ij}), \frac{\mathrm{Var}(S_{ij})}{n}\right) \sim \mathcal{N}\left(X, \frac{X(1-X)}{n}\right).$$

Since the dynamics of fCpG sites are by definition symmetric, the sample distribution of $Y_i$ for sites starting in the fully methylated state (right peak) is accordingly

$$Y_i | \; right \sim \mathcal{N}\left(1 - X, \frac{X(1-X)}{n}\right).$$

Finally, for fCpG sites that originated in the hemi-methylated state (middle peak), we note that their stationary state is centered at $X = 0.5$, that is

$$Y_i | \; middle = \mathcal{N}\left(0.5, \frac{0.25}{n}\right).$$

Inserting the three peak components into the mixture model (1), we then obtain a 4-parameter model

$$Y_i \sim \psi_1 \mathcal{N}(X, X(1-X)/n) + \psi_2 \mathcal{N}(0.5, 0.25/n) + \psi_3 \mathcal{N}(1 - X, X(1-X)/n). \tag{2}$$
