## Supplementary Figures for "Aggressive Neuroblastomas Start Growing after Infancy"

Monyak et al., 2026

### Content

**Supplementary Figure S1.** Selection of fluctuating CpG (fCpG) sites

**Supplementary Figure S2.** Tumor age vs. clinicopathological variables in the validation cohort.

**Supplementary Figure S3.** Pathway enrichment analysis.

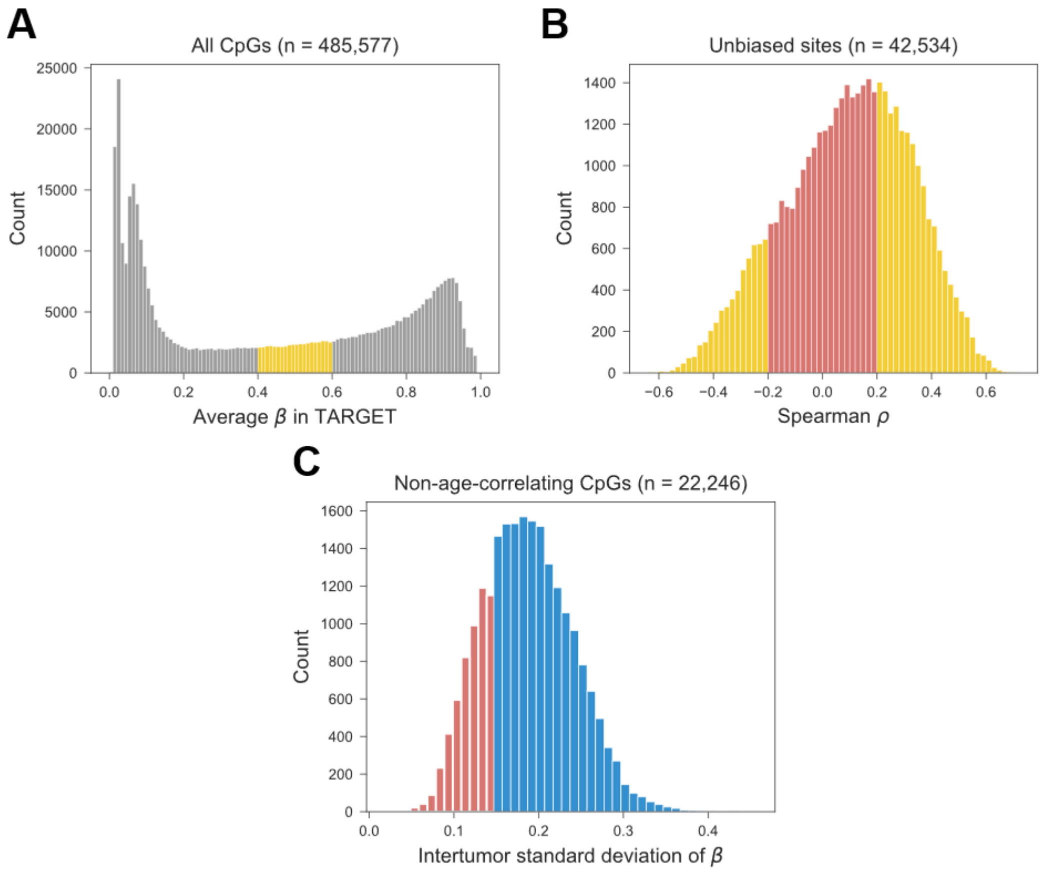


**Supplementary Figure S1. Selection of fluctuating CpG (fCpG) sites. (A)** Starting with all CpG sites (gray), *Unbiased Sites* were identified as those with an average methylation value $\beta$ between 0.4 and 0.6 in neuroblastomas (N = 213, TARGET cohort). **(B)** Starting with the set of *unbiased sites*, the spearman correlation between the methylation value $\beta$ and the patient age at diagnosis was calculated for each site, and sites with $|\rho| < 0.2$ (methylation uncorrelated with age) were selected for further refinement. **(C)** Non-age-correlating sites were ranked according to the inter-tumor standard deviation of their $\beta$-values, and the bottom 25% sites (red) were eliminated from further consideration.

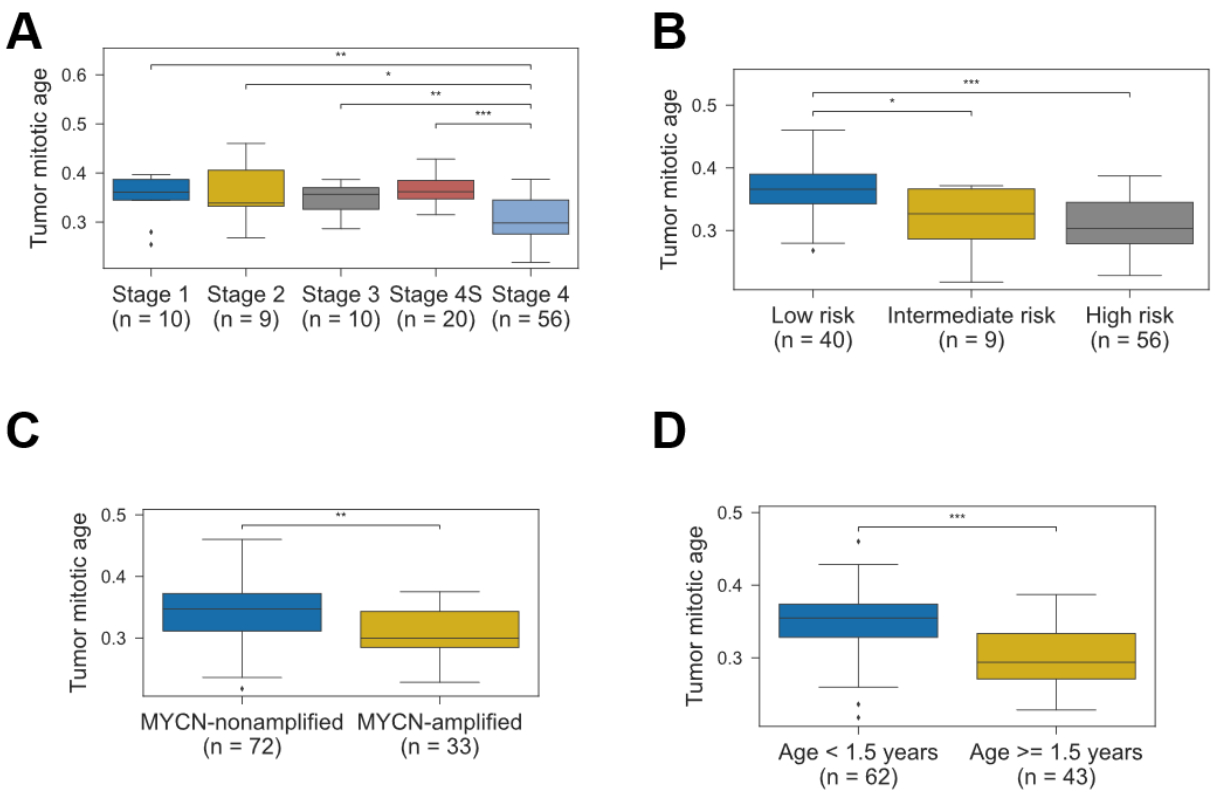


**Supplementary Figure S2. Tumor age vs. clinicopathological variables in the validation cohort**. The distribution of tumor age values among NBs in the validation cohort (N = 105), by **(A)** tumor stage; **(B)** risk category; **(C)** MYCN amplification status**;** and **(D)** patient age stratified at 1.5 years. Pairwise comparisons of medians were performed using a two-sided Wilcoxon rank-sum test (*P< 0.05, **P< 0.01, ***P< 0.001).

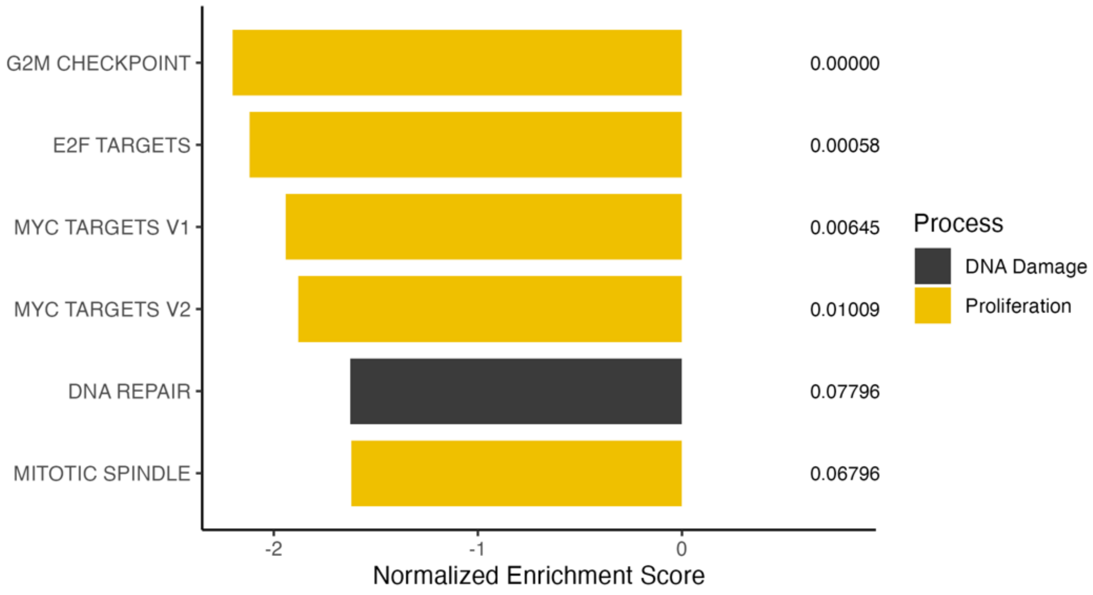


**Supplementary Figure S3. Pathway enrichment analysis.** A gene set enrichment analysis (GSEA) was performed for tumor mitotic age in neuroblastomas in the TARGET cohort with gene expression data (n = 132). The only significant associations found were negative associations between tumor mitotic age and Proliferation or DNA Damage-related pathways (genes enriched in mitotically younger tumors). Significant pathways were those with a false discovery rate (FDR) below 0.1 are shown; *FDR < .05; **FDR < .01; ***FDR < .001; ***FDR < .0001.
